# Contribution of the β-glucosidase BglC to the Onset of the Pathogenic Lifestyle of *Streptomyces scabies*

**DOI:** 10.1101/192591

**Authors:** Samuel Jourdan, Isolde M. Francis, Benoit Deflandre, Elodie Tenconi, Jennifer Riley, Sören Planckaert, Pierre Tocquin, Loïc Martinet, Bart Devreese, Rosemary Loria, Sébastien Rigali

## Abstract

Common scab disease on root and tuber plants is caused by *Streptomyces scabies* and related species which use the cellulose synthase inhibitor thaxtomin A as main phytotoxin. Thaxtomin production is primarily triggered by the import of cello-oligosaccharides. Once inside the cell, the fate of the cello-oligosaccharides is dichotomized into i) fueling glycolysis with glucose for the saprophytic lifestyle through the action of β-glucosidase(s) (BG), and ii) eliciting the pathogenic lifestyle by inhibiting the CebR-mediated transcriptional repression of thaxtomin biosynthetic genes. Here we investigated the role of *scab57721* encoding a putative BG (BglC) in the onset of the pathogenicity of *S. scabies*. Enzymatic assays showed that BglC was able to release glucose from cellobiose, cellotriose and all other cello-oligosaccharides tested. Its inactivation resulted in a phenotype opposite to what was expected as we monitored reduced production of thaxtomin when the mutant was cultivated on media containing cello-oligosaccharides as unique carbon source. This unexpected phenotype could be attributed to the highly increased activity of alternative intracellular BGs, probably as a compensation of *bglC* inactivation, which then prevented cellobiose and cellotriose accumulation to reduce the activity of CebR. In contrast, when the *bglC* null mutant was cultivated on media devoid of cello-oligosaccharides it instead constitutively produced thaxtomin. This observed hypervirulent phenotype does not fit with the proposed model of the cello-oligosaccharide-mediated induction of thaxtomin production and suggests that the role of BglC in the route to the pathogenic lifestyle of *S. scabies* is more complex than currently presented.

## Introduction

*Streptomyces scabies* is the causative agent of common scab on tuber and root plants via the production of the phytotoxin thaxtomin A amongst other virulence factors (Bignell *et al.*, 2010; Lerat *et al.*, 2009; Loria *et al.*, 2008). The onset of thaxtomin A is triggered upon transport of the cello-oligosaccharides cellobiose [(Glc)_2_] and cellotriose [(Glc)_3_] which involves the ATP-binding cassette (ABC) transporter system CebEFG-MsiK (Jourdan *et al.*, 2016). Once inside the cell, mainly cellobiose (Glc)_2_ but also cellotriose (Glc)_3_ can interact with the cellulose utilization repressor CebR preventing it from binding to its operator sequences associated with the thaxtomin biosynthetic gene cluster and therefore allowing the production of the phytotoxin (Francis *et al.*, 2015). Adjacent to the *cebR*-*cebEFG* divergon and 146 nucleotides downstream of *cebG*, *scab57721* (*bglC*) encodes a putative β-glucosidase (BG) of the glycosyl hydrolases (GH) GH1 family which is expected to catalyze the hydrolysis of terminal, non-reducing β-D-glucosyl residues, with release of β-D-glucose from β-D-glucosides and oligosaccharides (Henrissat, 1991; ENZYME entry: EC 3.2.1.21). The presence of a gene coding for an intracellular GH within the cluster of a sugar ABC-transporter is a common feature which allows co-transcription of genes required for carbohydrate import and their subsequent enzymatic degradation in the cytoplasm. Using molecules that are also common - most likely the most recurrent - soil carbohydrate nutrients for the onset of pathogenicity is very intriguing (Jourdan *et al.*, 2017). In non-pathogenic *Streptomyces*, coordinated expression of genes for BG and cello-oligosaccharide transport is appropriate for feeding the glycolysis pathway with glucose (Fig. 1). However, as stated earlier, in the plant pathogen *S. scabies*, (Glc)_2_ and (Glc)_3_ are not only perceived as nutrients used in the course of saprophytic behavior, but are above all signaling molecules eliciting its pathogenic lifestyle (Johnson *et al.*, 2007; Jourdan *et al.*, 2016; Wach *et al.*, 2007). Enzymes with a BG activity could thus potentially play an important role in controlling the onset of the virulence of *S. scabies* by limiting the intracellular accumulation of signals triggering thaxtomin A biosynthesis (Fig. 1). As a consequence, intracellular BG(s) of *S*. *scabies* might have evolved to display specific/unique properties which would ensure the microorganism to adopt the proper behavior – saprophytic versus pathogenic – according to environmental conditions (Fig. 1). In this work we defined the enzymatic properties, assessed the expression control mechanism, and investigated the role of *scab57721* (*bglC*) in thaxtomin A production and therefore in the onset of the virulence of *S. scabies.*

**Figure 1.**
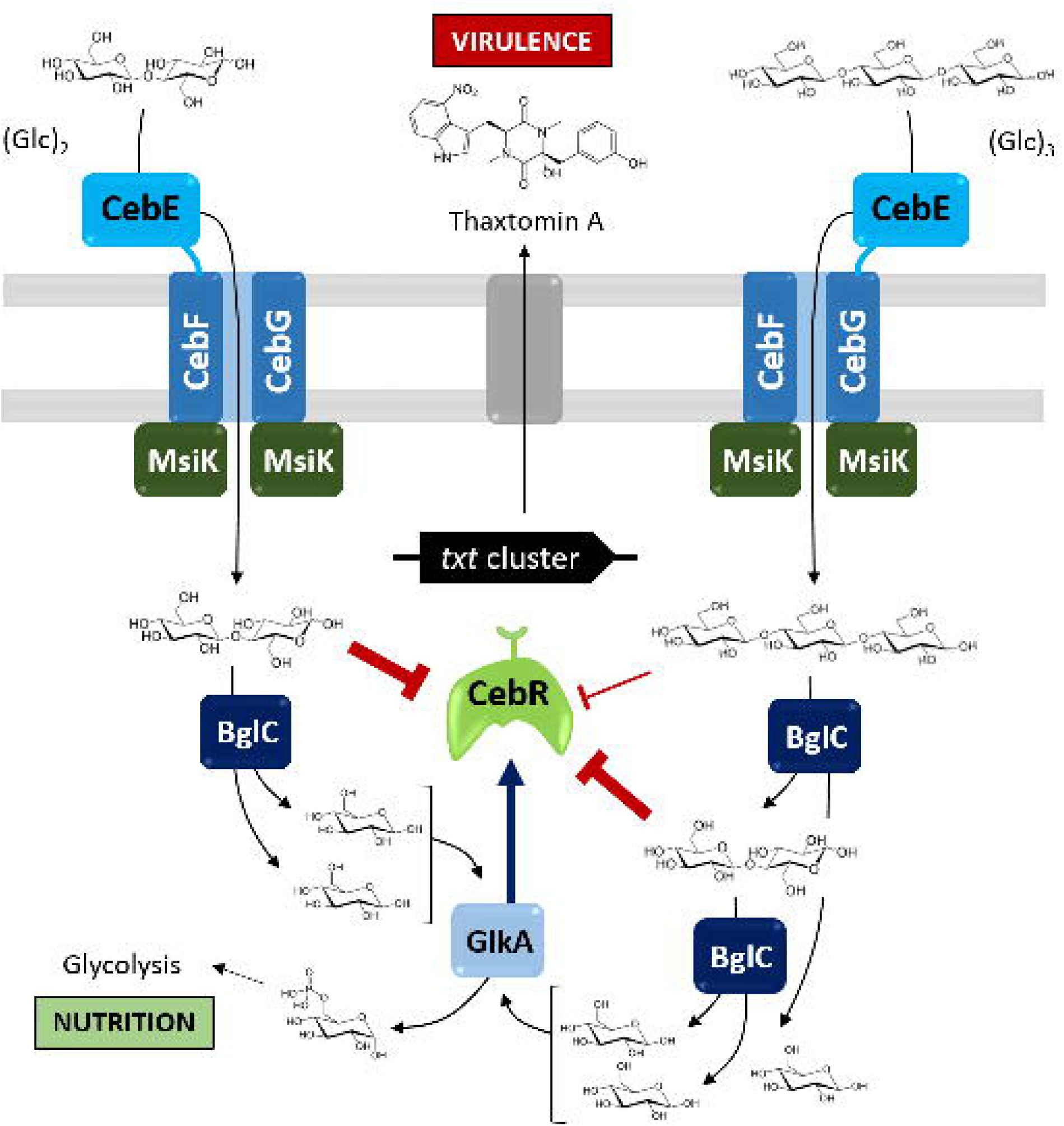
Position of the β-glucosidase activity on the modeled metabolic pathways from cellobiose and cellotriose transport to glycolysis and thaxtomin A production. When cellobiose and cellotriose are transported into the cytoplasm through the CebEFG-MsiK transporter, they both prevent the DNA-binding ability of the repressor CebR (cellobiose does this much more efficiently than cellotriose; Francis *et al.*, 2015), thus allowing expression of CebR-controlled genes including the thaxtomin biosynthetic genes (*txt* cluster), *cebEFG* and *bglC*. Once expressed, BglC cleaves both the imported cellobiose and cellotriose. Cellobiose hydrolysis directly leads to two glucose molecules, while cellotriose hydrolysis generates first glucose and cellobiose, the latter being the best allosteric effector of CebR. Cellotriose uptake would therefore inhibit CebR-mediated repression better than cellobiose uptake. The glucose generated by the BglC activity will be further metabolized by entering the glycolysis with phosphorylation by the glucose kinase GlkA to glucose-6-phosphate as the first step.

## Results and Discussion

### Enzymatic properties of BglC of *S. scabies*

The gene s*cab57721* encodes a 480 amino acid peptide orthologous to the well-characterized intracellular GH1 family BG BglC of *Thermobifida fusca* (53 and 67 % of amino acid identity and similarity, respectively) which also lies downstream of the *cebEFG* operon (Spiridonov and Wilson, 2001). BglC of *S. scabies* contains the MYVTENGAA sequence (amino acids 376 to 384) which matches the GH1 family active site signature [LIVMFSTC]-[LIVFYS]-[LIV]-[LIVMST]-E-N-G-[LIVMFAR]-[CSAGN] (PROSITE accession number PS00572). In order to assess the substrate specificity and the enzymatic properties of the predicted intracellular BG, s*cab57721* (*bglC*) was cloned into pET-28a (Table 1) for heterologous expression in *Escherichia coli* with a six histidine-tag fused to the N-terminus part of the protein (6His-BglC). Purification through Ni-NTA affinity chromatography enabled the recovery of 6His-BglC with an apparent molecular weight (MW) of ∼54 kDa which corresponds well to the its calculated MW of 54.121 kDa (Fig. 2A).

**Table 1.**
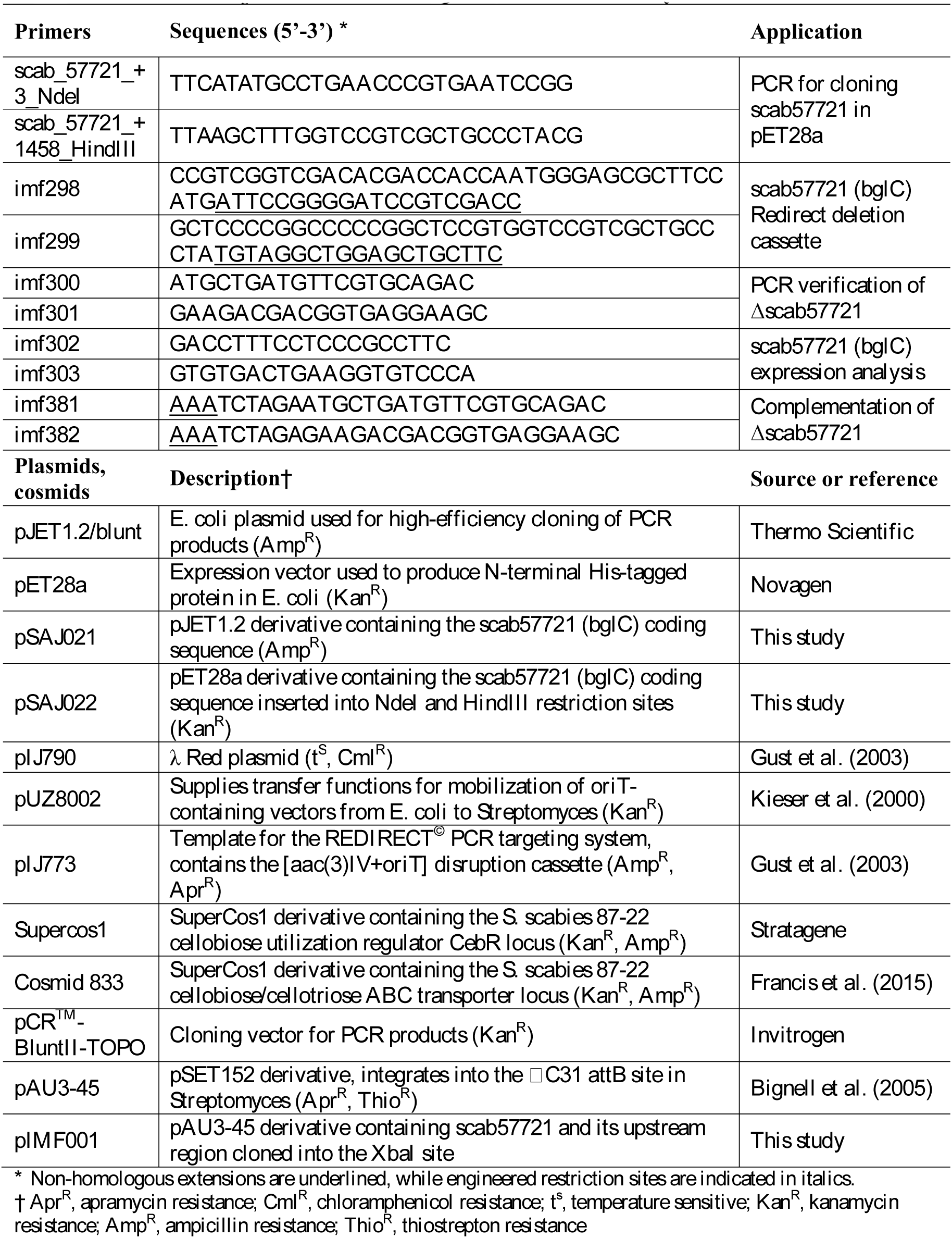
Primers and plasmids used and generated in this study.

**Figure 2.**
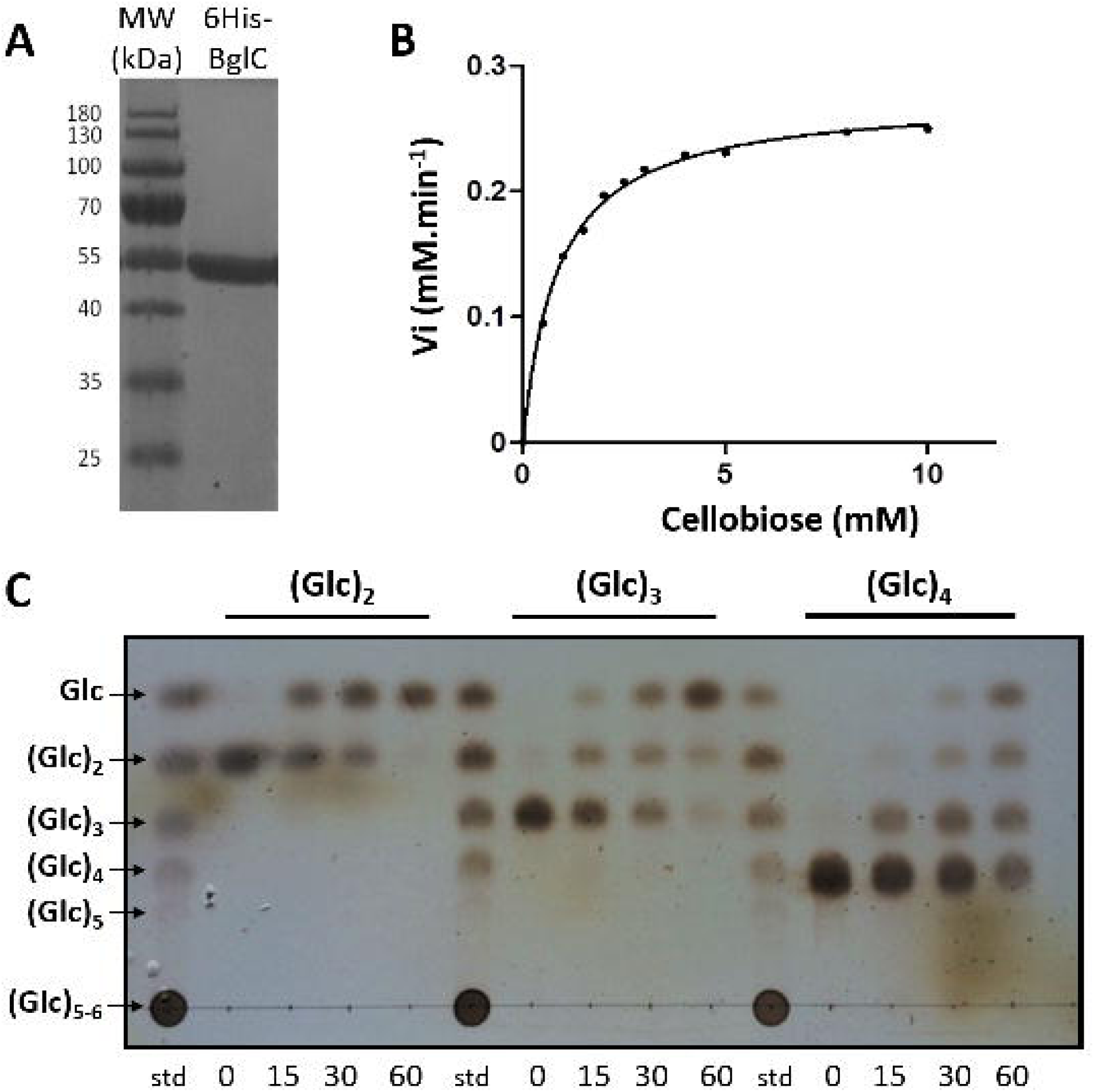
*scab57721* encodes a β-glucosidase. (A) SDS-PAGE showing the level of purity of 6His-BglC used for enzymatic assays. Lane 1, molecular weight marker; Lane 2, purified 6His-BglC of which the migration size (54 kDa) corresponds well to its predicted calculated size (54.121 kDa). (B) Initial velocity (V_i_) of 6His-BglC in function of the cellobiose concentration. Rates of cellobiose degradation were obtained by measuring the glucose released at the beginning of the hydrolysis reaction performed in 50 mM HEPES buffer pH 7.5 at 25°C. Data were fitted to the Henri-Michaelis-Menten equation using the GraphPad Prism 5 software in order obtain *V*_max_, *K*_*m*_, and *k*_*cat*_. (C) Substrate specificity of 6His-BglC for cello-oligosaccharides. Cello-oligosaccharides (6.25 mM) were incubated with pure 6His-BglC (0.4 μM) at 30°C for 0, 15, 30 and 60 min. std, standard cello-oligosaccharides: Glc, glucose; (Glc)_2_, cellobiose; (Glc)_3_, cellotriose; (Glc)_4_, cellotetraose.

The kinetic parameters of 6His-BglC were determined by measuring the initial rate of cellobiose hydrolysis (glucose release) at various concentrations of cellobiose. The maximum rate of the reaction (Vmax) is 7.3 μmol min^-1^ mg^-1^. The K_m_ and k_cat_ values were 0.77 mM and 400 min^-1^, respectively (Fig. 2B). The activity of 6His-BglC at different temperatures (from 20 to 55°C) and pH (from 5 to 10) values was measured using *p*-nitrophenyl-β-D-glucopyranoside (*p*-NPβG) as substrate (mimicking cellobiose). The activity of the enzyme gradually increased from 20 to 30°C, remained constant up to 37°C, and declined abruptly to 10% of the maximal activity at 42°C (Fig. S1). The optimal pH of BglC is around 7.5 as the enzyme maintained high activity between pH 6.5 and 8.5, and declined rapidly to 30 and 50% of its optimum at pH 5.5 and 9, respectively (Fig. S1).

To determine the substrate specificity of BglC, the recombinant protein was incubated with cellobiose (Glc)_2_, various cello-oligosaccharides ranging from cellotriose (Glc)_3_ to cellohexaose (Glc)_6_, as well as with different disaccharides unrelated to cellulose degradation (lactose, saccharose, maltose, threhalose, and turanose). Samples collected after increasing incubation times were spotted on a thin layer chromatography plate and revealed that 6His-BglC was able to generate glucose from cellobiose and all other cello-oligosaccharides tested (Fig. 2C). 6His-BglC was not able to release glucose from disaccharides unrelated to cellulose except for lactose though with much lower efficiency compared to cellobiose or any of the other cello-oligosaccharides (data not shown). If BglC displayed activity *in vitro* against (Glc)_4_, it is unlikely to occur inside the cytoplasm as the extracellular ABC transporter component CebE of *S. scabies* only displayed a high binding affinity to (Glc)_2_ and (Glc)_3_ (Jourdan *et al.*, 2016).

### *bglC* expression is repressed by CebR and induced by cellobiose

In order to ascertain that BglC is indeed involved in the catabolism of (Glc)_2_ and (Glc)_3_ *in vivo*, we assessed if its expression/production in *S. scabies* is under the control of the cellulose utilization repressor CebR. Quantitative reverse transcription PCR (qPCR) was performed on RNA extracted from the wild-type strain of *S. scabies*, 87-22, and its *cebR* deletion mutant, Δ*cebR*, grown on ISP-4. This revealed that the deletion of *cebR* resulted in an 85-fold (wild-type 0.025 vs. *cebR* mutant 2.13) overexpression of *bglC* (Fig. 3A). In addition, targeted LC-MRM analysis allowed evaluation of the effect of the deletion of *cebR* as well as the presence of cellobiose on BglC production in *S. scabies.* Quantitative analyses of two specific tryptic peptides of BglC (LVDELLAK and TDPVASLR) showed that the protein was more abundant in the total intracellular protein extracts of the Δ*cebR* mutant (2.3 fold more compared to 87-22) as well as in extracts of the *S. scabies* wild-type strain grown in cellobiose-containing media (2.9 fold more compared to the condition without cellobiose) (Fig. 3B). The observed transcriptional repression exerted by CebR and the cellobiose-dependent induction of *bglC*/BglC are mediated through direct binding of CebR to the CebR-binding site (TGGaAGCGCTCCCA) identified at position -14 nt upstream of *bglC* (Fig. 3C). The results deduced from the targeted proteomic approach are in agreement with the early and constitutive overall intracellular BG activity measured as a consequence of *cebR* deletion while *S. scabies* 87-22 wild-type only displayed measurable BG activity when grown in the presence of cellobiose (Fig. 3D).

**Figure 3.**
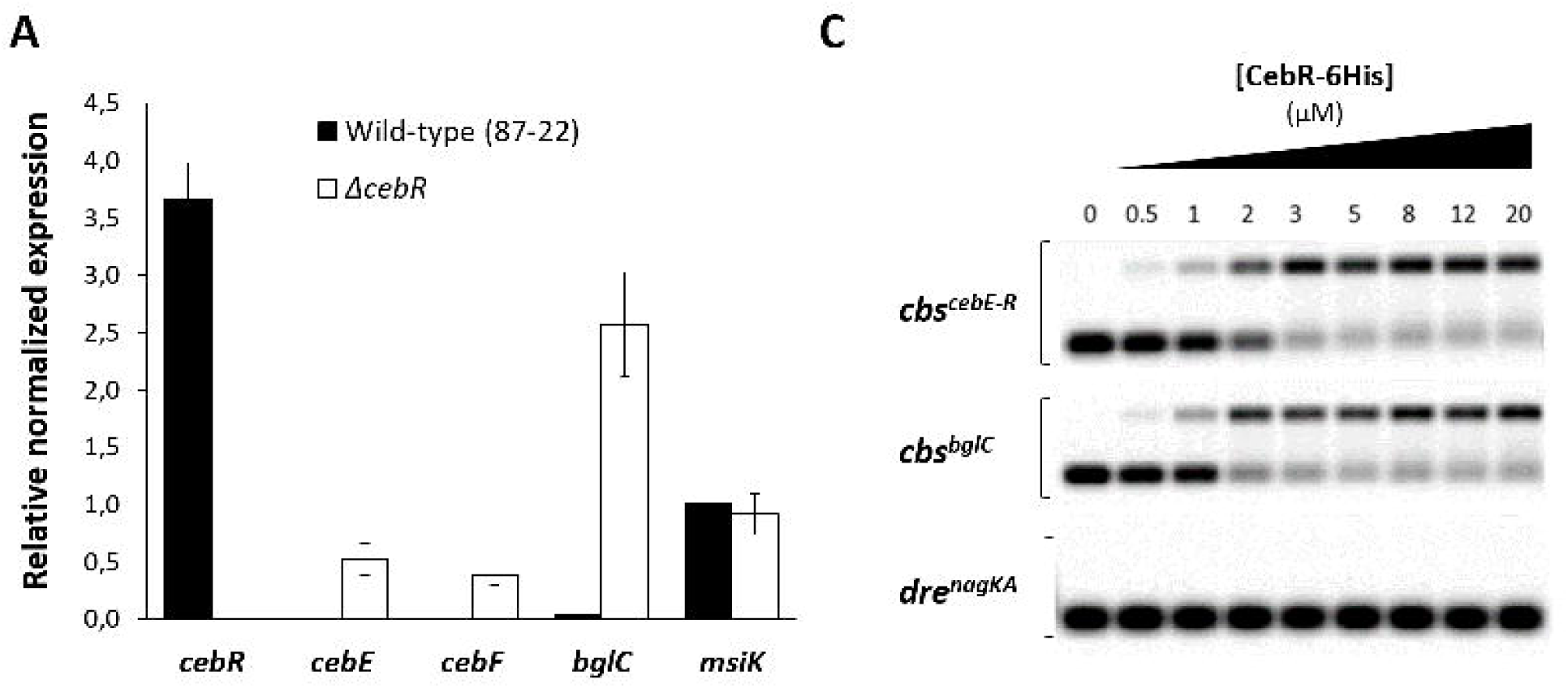
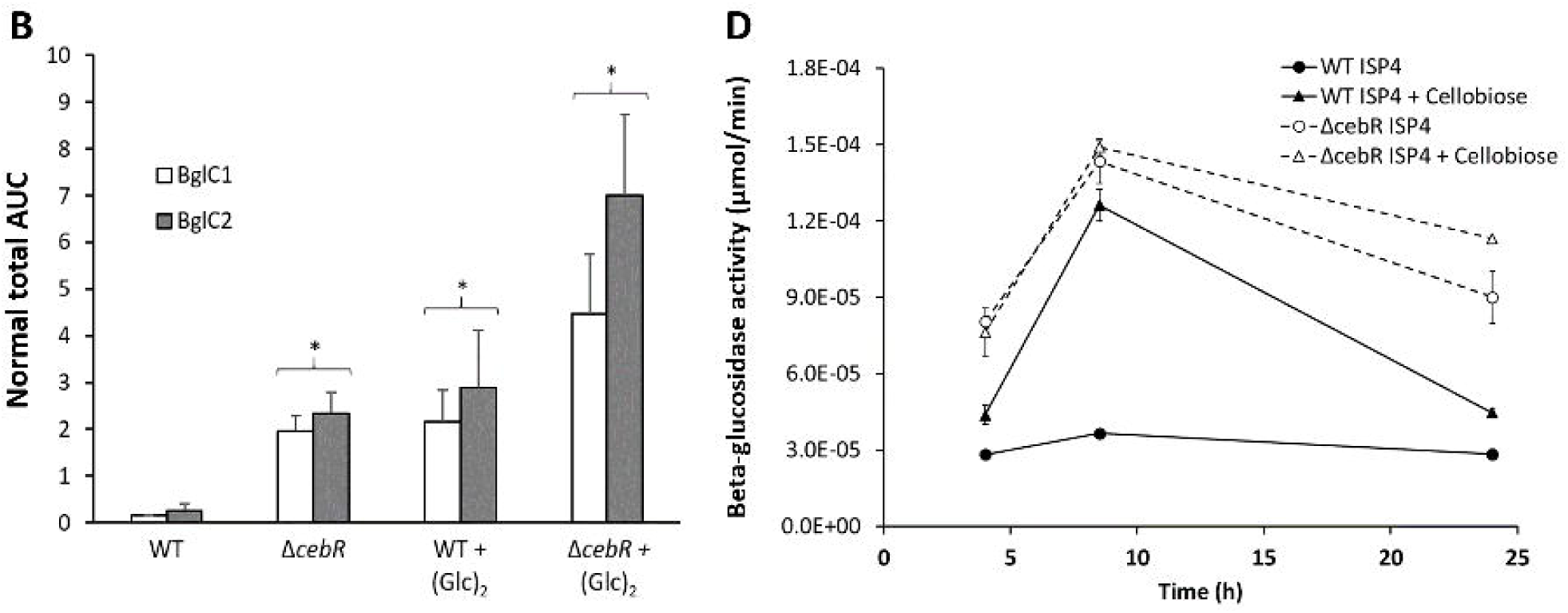
Expression of *bglC* is repressed by CebR and induced by cellobiose. (A) qPCR analysis of *bglC* expression levels in *S. scabies* 87–22 and in the Δ*cebR* strain. Data were normalized using the *gyrA* and *murX* genes as internal controls and using *cebE*, *cebF*, and *cebR* as CebR repressed genes. Mean normalized expression levels (± standard deviations) from three biological repeats analyzed in triplicate are shown. (B) Relative normalized abundancy of BglC peptides in response to the deletion of *cebR* (Δ*cebR*) and/or cellobiose supply, determined by LC-MRM MS on tryptic digests of protein extracts. Target peptides for BglC: LVDELLAK (BglC1) and TDPVASLR (BglC2). * denotes significant quantitative peptide overproduction (p < 0.05) compared to the wild-type (WT) strain grown in ISP-4 without cellobiose supply. Statistical significance was assigned by performing 2-sided Student’s t-tests and assuming groups of equal variances. (C) EMSAs showing specific interaction of CebR with the *cbs* (CebR-binding site) element at position -14 nt upstream of *bglC*. Probes with the DasR-responsive element (*dre*) upstream of *nagKA* (Tenconi *et al.*, 2015) and with the *cbs* upstream of *cebE* were used as negative and positive controls, respectively. (D) Overall β-glucosidase activity of *S. scabies* 87-22 and its *bglC* null mutant grown in liquid ISP4 with or without cellobiose (0.5 mM) supply.

### Inactivation of *bglC* results in reduced thaxtomin A production when *S. scabies* is grown with cello-oligosaccharides as the sole carbon source

Since we demonstrated that *bglC*/BglC i) is induced by cello-oligosaccharides, and ii) displays BG activity against (Glc)_2_ and (Glc)_3_, we finally assessed if the catabolic activity of BglC influenced the production levels of thaxtomin A and as a consequence virulence of *S*. *scabies*. We generated a *bglC* null-mutant (Δ*bglC*) by replacing *orf scab57721* by the apramycin resistance cassette as performed previously for *cebR*, *cebE* and *msiK* (Francis *et al.*, 2015; Jourdan *et al.*, 2016). Semi-quantitative analysis by HPLC revealed that the Δ*bglC*/Δ*57721* mutant under-produced thaxtomin A to only 37% and 9% of the thaxtomin levels produced by the wild-type strain when cultivated in liquid minimal medium (MM) with cellobiose or cellotriose as sole carbon sources (Fig. 4A). This result was unexpected as the deletion of *bglC* should normally lower the catabolism of cello-oligosaccharides for glycolysis and therefore would result in their higher intracellular accumulation as allosteric inhibitors of CebR and activators of *txtR* expression.

**Figure 4.**
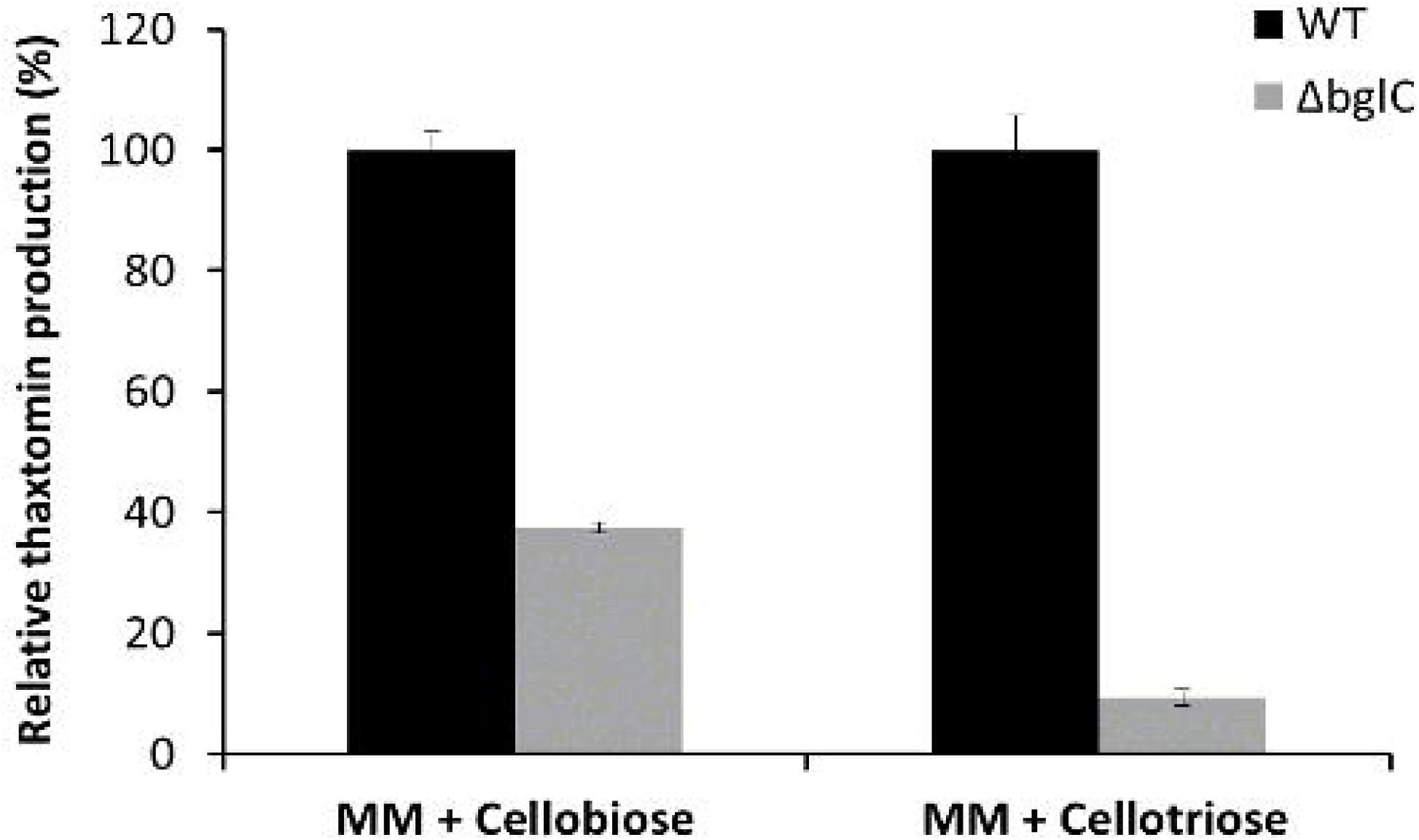
Effect of *bglC* deletion on the cello-oligosaccharide-mediated induction of thaxtomin A production. *S. scabies* 87-22 and its *bglC* null mutant were grown in liquid MM medium supplemented with 0.5 mM cellobiose or cellotriose. Thaxtomin production was quantified by HPLC after 24h post-inoculation and wild-type production levels in each condition were fixed to 100%.

To tentatively explain the reduced thaxtomin A production as a result of the inactivation of *bglC*, we monitored cellobiose or cellotriose consumption as well as the total BG activity of the Δ*bglC* strain. For this purpose, *S. scabies* wild-type 87-22 and its *bglC* null mutant were grown for 24 hours in MM supplemented with either 500 μM of (Glc)_2_ or (Glc)_3_ as the sole carbon source. The concentration of cello-oligosaccharides remaining in the culture supernatant was measured by HPLC at 1.5 hour intervals post inoculation (hpi) (Fig. 5A). Full consumption of cellobiose and cellotriose by the wild-type strain 87-22 was accomplished at 3 and 4.5 hpi, respectively, while the *bglC* mutant was impaired in both cellobiose and cellotriose utilization as total consumption of these cello-oligosaccharides required about 3 h longer than for the wild-type (Fig. 5A). This delayed import and consumption could possibly postpone the production of thaxtomin A in the *bglC* mutant but should not be responsible for the observed massive reduction of production of the phytotoxin.

**Figure 5.**
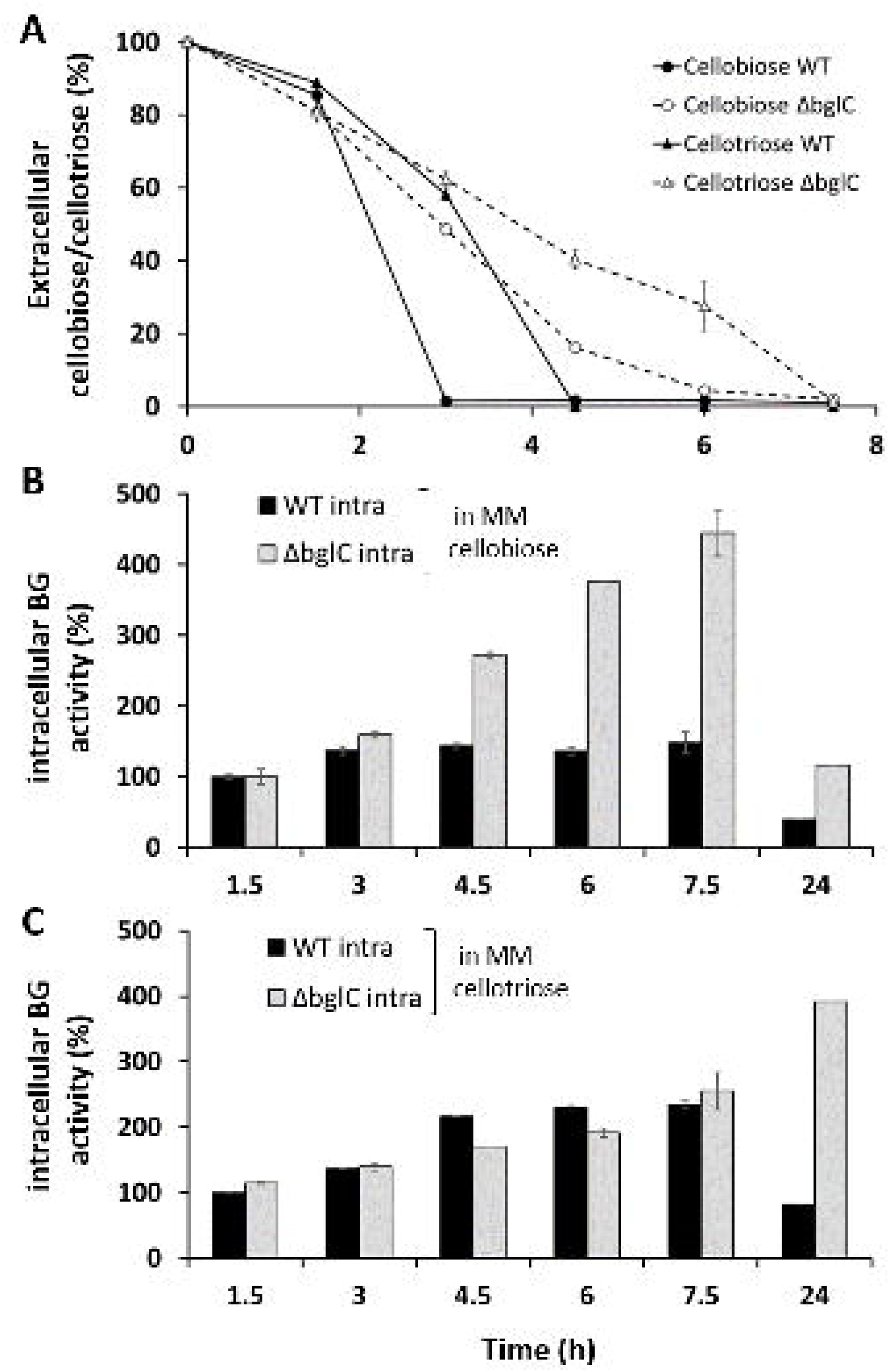
Consumption of the cello-oligosaccharides cellobiose and cellotriose (A), and correlation with the intracellular β-glucosidase activity of *S. scabies* wild-type and the *bglC* null mutant (B) and (C). The BG activity of the wild-type at the first time point was set to 100%.

Concomitantly to the measurements of the cello-oligosaccharide consumption, we assessed the intracellular and extracellular relative BG activity to evaluate to which extent the loss of *bglC* impacted the overall BG activity (Figs. 5BC and S2). Each soluble fraction (intra-and extracellular) was assessed at five different time points in both wild-type and Δ*bglC* strains using *p-*nitrophenyl-β-D-glucopyranoside (*p*-NPβG) as substrate. Very low extracellular BG activities were obtained for both strains and under both MM with cellobiose or cellotriose culture conditions (Fig. S2). Assessment of the intracellular BG activity against *p*-NPβG revealed that while the activities measured in *S. scabies* wild-type and Δ*bglC* were similar in the cellobiose-containing medium at the beginning of the culture, the activity of the mutant strain increased dramatically after 3 hpi (Fig. 5B). At the end of the experiment, the wild-type strain presented only a slight increase in BG activity reaching merely one third of the overall BG activity displayed by the *bglC* null mutant (Fig. 5B). The corresponding activities measured in cellotriose-containing medium were more similar for both strains at the beginning of the culture but the *bglC* null mutant presented a BG activity that was about 4 times higher than that of the WT at 24 hpi (Fig. 5C). This delay in the response of the BG might be a consequence of the delay in cellotriose consumption observed for the Δ*bglC* strain (Fig. 5A) but also because cellotriose is a much weaker allosteric effector of CebR compared to cellobiose (Francis *et al.*, 2015). These observations demonstrate that BglC is not the only functional β-glucosidase in *S. scabies* to catabolize cello-oligosaccharides. The fact that the mutant displayed BG activity points to the presence of one or several additional/alternative β-glucosidases which are apparently overproduced or for which the biosynthesis is awakened when cellobiose or cellotriose was provided as the sole carbon source. The nature and the pathway associated with the induction of the alternative β-glucosidase(s) are currently unknown but might involve CebR as the response differed according to cellobiose or cellotriose supply. The contribution of BglC to the overall BG activity of the wild-type is another pending question.

That the *bglC* null mutant displayed a much higher overall BG activity would result in a more rapid depletion of the incorporated thaxtomin-inducing cello-oligosaccharides, thus providing a possible explanation of the unexpected decreased thaxtomin A production of *S*. *scabies* Δ*bglC* compared to the wild-type when cello-oligosaccharides are provided as the only carbon source. Similar reduced thaxtomin A production levels were also observed when assays were performed on solid MM. When inoculated on MM with cellobiose as sole carbon source (TDMc, Fig. 6), the Δ*bglC* mutant displayed a growth delay during the first 24h consistent with the absence of a major cellobiose hydrolyzing enzyme. When incubated for a longer period, growth is recovered but the Δ*bglC* strain cannot reach the level of thaxtomin produced by the wild-type in TDMc as previously described in liquid minimal medium (Fig. 4A).

**Figure 6.**
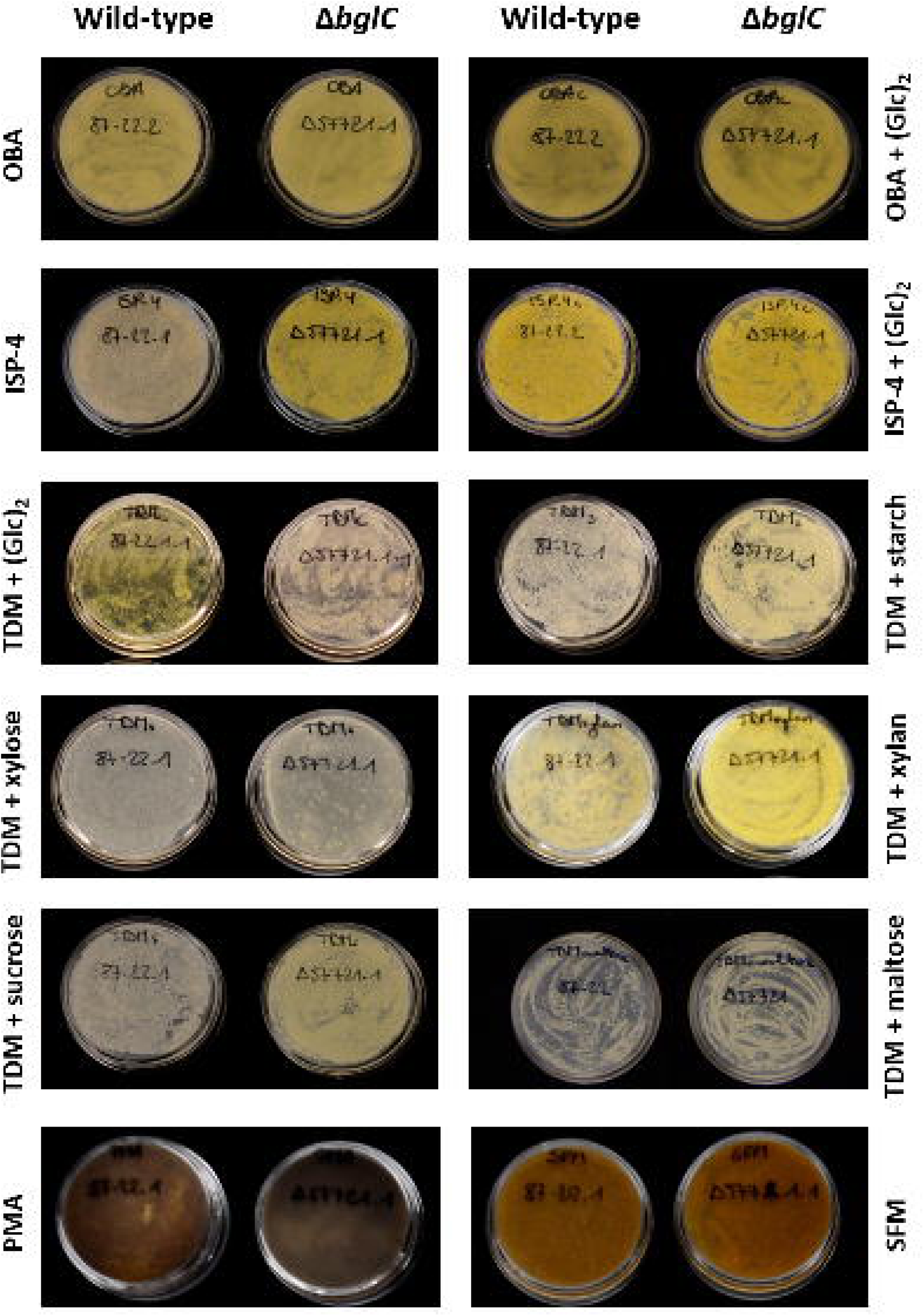
Thaxtomin A production by *S. scabies* wild-type (87-22) and the *bglC* null mutant grown on various minimal and complex solid media. (A) Pictures of media inoculated with *S. scabies* 87-22 and its *bglC* null mutant. Thaxtomin A production can be seen due to its distinct yellow pigmentation. (B) Quantification of thaxtomin A extracted from plates shown in A after incubation for 7 days at 28°C. Means and standard deviations were calculated on three biological replicates. The wild-type production level in TDM cellobiose was fixed to 100%.

### Inactivation of *bglC* results in overproduction or constitutive production of thaxtomin A when cell-oligosaccharides are not the only carbon source

The capability of the mutant to produce thaxtomin A was also monitored on a series of solid media amongst which the complex OBA medium that naturally contains cello-oligosaccharides and other carbon sources (Johnson *et al.*, 2007; Fig. 6). When grown on OBA the Δ*bglC* mutant overproduced thaxtomin A compared to the wild-type strain (Fig. 6B). On this medium, the addition of cellobiose to the OBA medium neither decreased nor further increased thaxtomin production suggesting that the *bglC* mutant could have partially lost its capacity to respond to cellobiose when other carbon sources are available (Fig. 6B). Surprisingly, the *bglC* mutant also overproduced thaxtomin A when inoculated on ISP4 medium deprived of cello-oligosaccharides as nutrient sources (Fig. 6). In order to ascertain the validity of this unexpected phenotype, the mutant was complemented by introducing plasmid pIMF001 (Table 1) containing the *bglC* gene with its promoter into the Δ*bglC* mutant isolates. Complementation of Δ*bglC* restored the wild-type phenotype when bacteria were streaked out on ISP-4 (Fig. S3) demonstrating that the observed alteration in thaxtomin production was indeed caused by the deletion of the *bglC* (*scab_57721*) gene and not due to a possible unspecific event such as a spontaneous mutation. The thaxtomin A overproduction phenotype was further confirmed on most media tested, so regardless of the presence of cellobiose or other cello-oligosaccharides (Figure 6).

Since Δ*bglC* showed constitutive production of thaxtomin A, its virulence capacity was evaluated on *Arabidopsis thaliana* and radish seedlings. No different outcome was observed between radish seedlings infected with the wild-type or the mutant (Fig. 7A). However, since the outcome of the radish assay is mostly influenced by the effect of thaxtomin on the plant’s growth and development and thaxtomin is active in nanomolar concentrations (King *et al.*, 2001), it is hard to see any difference between the production levels of the wild-type and a potential thaxtomin overproducer using radish as host. Assays were also performed using slightly older seedlings (48h instead of 30h after sowing) or a lower inoculum (200 μl of a mycelial stock of OD_600_ 0.1 instead of OD_600_ 1.0), still no difference could be observed. Yet, when thaxtomin was extracted from the agar-water support with the radish seedlings, a significantly higher concentration of thaxtomin A was measured for the assays done with the Δ*bglC* isolates compared to the wild-type strain (Fig. S4). The use of *A. thaliana* (ecotype Col-0) as the plant model revealed to be more suitable for monitoring hypervirulent phenotypes than radish seedlings as previously observed for the *cebR* mutant which also overproduces thaxtomin A (Francis *et al.*, 2015). *A. thaliana* seeds grown on Murashige-Skoog (MS) agar were inoculated with spores of *S. scabies* 87-22 (wild-type) and its Δ*bglC* mutant. After 7 days of growth, seedlings inoculated with the *bglC* mutant presented stronger growth and developmental defects compared to those inoculated with the wild-type strain (Fig. 7B). Closer inspection of individual plants revealed stronger root and shoot stunting as a consequence of the *bglC* deletion (Fig. 7B).

**Figure 7.**
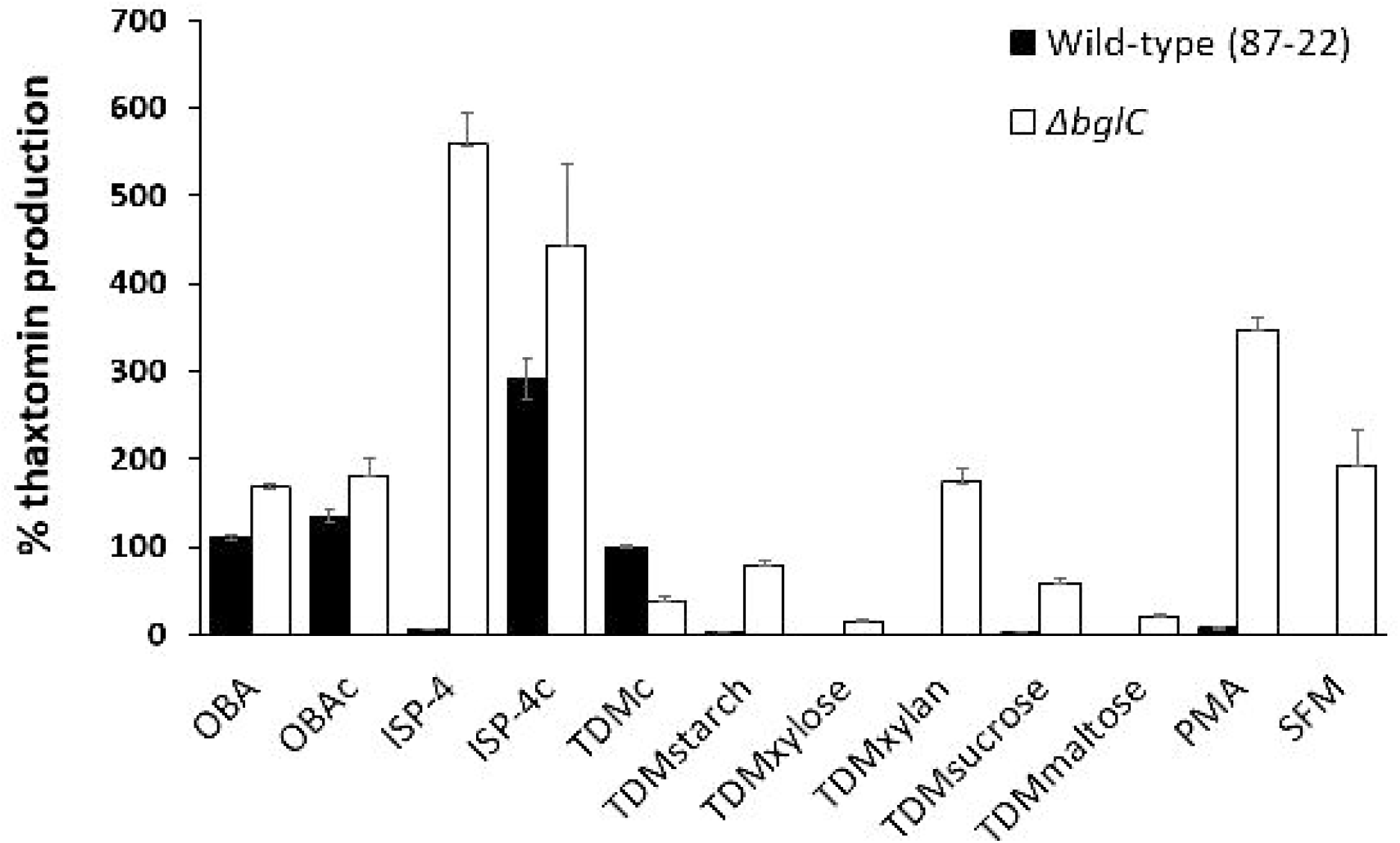
Effect of *bglC* deletion on the virulence of *S. scabies*. (A) Phenotypes of representative radish seedlings treated with water, the wild-type strain 87-22, and *bglC* mutant isolates at 6 days post infection. (B) Phenotype of *A. thaliana* grown for 7 days in the presence of *S. scabies* 87-22 (wild-type) and its *bglC* null mutant with a close-up of representative plants grown on the MS plates shown in the upper panel.

## Conclusion and perspectives

In this work we demonstrated that the protein encoded by the gene *scab57721* located downstream of the *cebEFG* operon is a β-glucosidase active against different cello-oligosaccharides including the best inducers of thaxtomin A production *i*.*e*., cellobiose and cellotriose. Expression of *bglC* is also repressed by CebR, the master regulator of pathogenicity in *S. scabies,* and induced by cellobiose. Since cellobiose and cellotriose consumption by *S. scabies* correlates with an intracellular increase of β-glucosidase activity, we assumed that BglC (and any other enzyme with BG activity) would play an essential role in controlling the pool of imported elicitors to trigger the CebR regulon and therefore thaxtomin production as proposed in the model illustrated in Fig. 1. In line with the current model of the cello-oligosaccharide-mediated induction of thaxtomin A production we were expecting that the inactivation of *bglC* would simply result in an increased or prolonged production of thaxtomin under culture conditions supplemented with cellobiose or cellotriose as these CebR-allosteric molecules would remain longer in the cytoplasm. However, surprisingly, we observed that the presence of cellobiose and cellotriose as sole carbon source instead reduced the production levels of thaxtomin A, probably as a consequence of the awakening of alternative BG(s) encoded in the genome of *S*. *scabies* as compensation for the loss of BglC. Identification of the protein(s) responsible for the high BG activity in the *bglC* mutant is currently under investigation.

Finally, the most striking phenotype observed for the Δ*bglC* strain was the loss of the cellobiose-dependent induction of thaxtomin and thus the constitutive thaxtomin production in complex media devoid of eliciting cellulose-related sugars (Fig. 6). That this mutant is able to produce thaxtomin without the presence of the inducing molecules is difficult to explain based on the current model of the induction pathway of thaxtomin production and suggests that the role of BglC in the induction of *S. scabies* pathogenicity involves mechanisms that still have to be uncovered.

## Experimental procedures

### Bacterial strains and culture conditions

*Escherichia coli* strains [DH5α and Rosetta^™^ (DE3)] were cultured in Luria-Bertani (LB) medium at 37°C. *Streptomyces* strains (wild-type 87-22 and mutant strains Δ*57721*/Δ*bglC*) were routinely grown at 28°C in tryptic soy broth (TSB; BD Biosciences) or on International *Streptomyces* Project medium 4 (ISP-4, BD Biosciences). When required, the medium was supplemented with the antibiotics apramycin (100 μg/ml), kanamycin (50 μg/ml), chloramphenicol (25 μg/ml), thiostrepton (25 μg/ml), and/or nalidixic acid (50 μg/ml). Cellobiose and cello-oligosaccharides were purchased from Megazyme (Ireland). For the BG activity assays and the thaxtomin production assays the *Streptomyces* strains were grown on the complex media Oat Bran Agar (OBA; Johnson *et al.*, 2007), Soy Flour Mannitol (SFM; (Kieser *et al.*, 2000), Potato Mash Agar (PMA; 12.5 g potato flakes and 5 g agar per liter), as well as the minimal medium Thaxtomin Defined Medium (TDM), modified from Johnson *et al.* (2007) by omitting xylose and using a final concentration of 1% of the carbon source of choice.

### Heterologous expression and purification of His-tagged BglC

The open reading frame encoding SCAB57721 (BglC) was amplified by PCR using the primers scab*_*57721_+3_*Nde*I and scab_57721_+1458_*Hind*III (see Table 1 for primer sequences). The PCR product was subsequently cloned into the pJET1.2/blunt cloning vector, yielding pSAJ021. After DNA sequencing to verify the correct amplification of *scab57721*, an *Nde*I-*Hind*III DNA fragment was excised from pSAJ021 and cloned into pET-28a digested with the same restriction enzymes leading to pSAJ022. *E. coli* Rosetta™ (DE3) cells carrying pSAJ022 were grown at 37°C in 250 ml LB medium containing 50 μg/ml of kanamycin until the culture reached an absorbance at 600 nm (A_600_) of 0.6. Production of 6His-tagged BglC (6His-BglC) was induced overnight (∼20 h) at 16°C by addition of 1 mM isopropyl-β-D-thiogalactopyranoside (IPTG). Cells were collected by centrifugation and ruptured by sonication in lysis buffer (100 mM Tris-HCl buffer; pH 7.5; NaCl 250 mM; 20 mM imidazole) supplemented with the EDTA-free cOmplete protease inhibitor cocktail (Roche). Soluble proteins were loaded onto a pre-equilibrated Ni^2+^-nitrilotriacetic acid (NTA)-agarose column (5-ml bed volume), and 6His-BglC was eluted within the range of 100 to 150 mM imidazole. Fractions containing the pure protein were pooled (Fig. 2A) and dialyzed overnight in 50 mM HEPES; pH 7.5.

### Construction of the *bglC* mutant in *S. scabies* 87-22 and its genetic complementation

The deletion of the *bglC* coding region was created as described previously (Francis *et al.*, 2015; Jourdan *et al.*, 2016). Specific primers used to generate and verify the gene deletion and complement the *bglC* null-mutant are listed in Table 1. A fragment containing the *bglC* coding region and the upstream region (379 bp) harboring the promoter was generated by PCR using primers with engineered *Xba*I sites (Table 1) and cloned into pCR^™^-BluntII-TOPO (Invitrogen). After sequence confirmation, fragments were retrieved through an *Xba*I restriction digest, gel purified, and cloned into an *Xba*I-linearized pAU3-45 (Bignell *et al.*, 2005) resulting in plasmid pIMF001 (Table 1). Complementation constructs, as well as the empty pAU3-45 plasmid, were introduced into three *bglC* mutant isolates through intergeneric conjugation similar to the gene deletion process as described previously (Francis *et al.*, 2015; Jourdan *et al.*, 2016).

### Quantitative Reverse Transcription PCR

RNA was prepared from 72-h-old mycelia grown on ISP-4 plates at 28°C using the RNeasy minikit (Qiagen) according to the manufacturer’s instructions. Verification of the absence of contaminating genomic DNA, cDNA synthesis, and quantitative reverse transcription PCR (qPCR) were performed as described previously (Francis et al. 2015; Jourdan et al. 2016). The *bglC* specific internal primers imf302 and imf303 were used to quantify the expression levels of the *bglC* gene (Table 1). The *murX*, *hrdB*, and *gyrA* genes were used to normalize the amount of RNA in the samples (Joshi *et al.*, 2007). Each measurement was performed in triplicate with three different *cebR* mutant isolates.

### Targeted proteomics

*S. scabies* 87-22 and its *cebR* null mutant were grown on ISP-4 plates with or without a 0.7% cellobiose supply. The mycelium was collected after 48 hours of incubation at 28°C, and resuspended in 50 mM NH_4_HCO_3_ buffer (pH 7.5). Crude intracellular extracts were obtained after sonication of the mycelium as described previously (Jourdan *et al.*, 2016). Sample preparation for Liquid Chromatography-Multiple Reaction Monitoring (LC-MRM) analysis, and LC-MRM analysis were performed as previously described (Jourdan *et al.*, 2016).

### β-glucosidase activity assays

The relative enzyme activity was determined using *p-*nitrophenyl-β-D-glucopyranoside (*p-* NPβG) as substrate. The reaction mixture (200 μl) containing 50 mM HEPES buffer (pH 7.5), 0.2 μM of purified 6His-BglC and the tested reagent was incubated for 10 min at 25°C before addition of 1 mM *p-*NPβG. The reaction was carried out at 25°C for 2 min and stopped by addition of 100 μl of 2 M Na_2_CO_3_. All assays were performed under these conditions, unless otherwise indicated. The release of *p*-nitrophenol (*p-*NP) was measured at 405 nm with a TECAN infinite^®^ 200 PRO.

### Kinetic analysis

Kinetic parameters of BglC (*K*_*m*_ and *k*_*cat*_) were determined by measuring the glucose released at various cellobiose concentrations in 50 mM HEPES buffer pH 7.5 at 26°C. A reaction time of 7 min was chosen to ensure initial rates of hydrolysis. The glucose released was determined using the D-Glucose HK Assay Kit from Megazyme (Ireland). Data were fitted to the Henri-Michaelis-Menten equation using the GraphPad Prism 5 software.

### Hydrolysis of disaccharides and oligosaccharides

The cleavage ability of BglC was tested against different cello-oligosaccharides (cellobiose, cellotriose, and cellotetraose, (Megazyme; Ireland) or different disaccharides (lactose, saccharose, maltose, threhalose and turanose). Reaction mixtures (100 μl) containing 50 mM HEPES buffer pH 7.5; 0.4 μM of purified 6His-BglC; 6.25 mM of cello-oligosaccharides or 12.5 mM of disaccharides were incubated at 30°C. Samples of each 15 μl were collected at 0, 15, 30 and 60 min, and heated at 98°C for 5 min to stop the reaction. Each sample was spotted onto an aluminum-backed Silica gel plate (Sigma). The plates were run with chloroform-methanol-acetic acid-water solvent (50:50:15:5, vol/vol), air dried, dipped in 5% H_2_SO_4_ in ethanol and heated over a hot plate until visualization of the carbohydrate spots as described by Gao and Wakarchuk (2014).

### Monitoring of cellobiose and cellotriose consumption and glucose production

Glucose, cellobiose, and cellotriose consumption measurements were performed by HPLC (Alliance, Waters Milford, MA, USA) on a lead-form Aminex HPX-87P Column (300 x 7.8 mm, 9μm particle size supplied by Bio-Rad) in combination with two Micro-Guard columns (De-Ashing refill cartridge 30 x 4.6 mm supplied by Bio-Rad) heated to 80°C with Milli-Q (18.2 MΩ cm) distilled-deionized H_2_O in an isocratic mode (flow rate 0.6 ml/min). Peaks were detected by a refractive index detector (Waters 2414) and processed with the Empower 3 software (Waters Milford, MA, USA).

### Thaxtomin production assays

Thaxtomin production assays were performed as described previously (Francis *et al.*, 2015; Jourdan *et al.*, 2016). Briefly, plates were inoculated with equal amounts of mycelial suspensions of the *S. scabies* 87-22 wild-type and its *bglC* null mutant, and incubated 7 days at 28°C. Thaxtomin was extracted from the agar and quantification by reversed-phase high-performance liquid chromatography (HPLC) was performed as described previously (Francis *et al.*, 2015; Jourdan *et al.*, 2016). For liquid cultures, thaxtomin was extracted from 1 ml of the culture supernatant with 0.3 ml of ethyl acetate and quantified by HPLC using a NUCLEODUR^®^ 100-5 C18ec column (Macherey-Nagel). Samples were eluted at a flow rate of 0.8 ml/min, and A400 was monitored using a Multi λ Fluorescence detector (2475, Waters). All experiments were repeated using three different biological and technological replicates per *S. scabies* strains.

### Virulence assays

Virulence assays on *Arabidopsis* seedlings were performed as follows. Seeds of Col-O ecotype were surface sterilized for 15 min in bleach solution (40% vol/vol bleach, 0.05% vol/vol Tween20), thoroughly rinsed with sterile H_2_O, and stratified 3 days at 4°C in the dark before sowing. 300 to 400 *Arabidopsis* seeds were sown in each well of a six-well plate containing half concentrated MS medium (Sigma M5513) supplemented with 1% sucrose. Each well was inoculated with 250 μl spore suspensions of the *S. scabies* 87-22 wild-type and *bglC* mutant (5.10^4^ spores per μl), or sterile water as the control. The plates were incubated at 25 ± 0.5°C under 16-h photoperiod for 7 days.

Virulence phenotypes on radish seedlings were performed as described previously (Jourdan *et al.*, 2016). Thaxtomin was extracted from the total of the radish seedlings and the water-agar medium by cutting the material into small pieces and soaking in 15 ml methanol for 10 min. The liquid phase was dried down and resuspended in 1 ml methanol. These samples were analyzed by HPLC as described above.

## Acknowledgments

S.J. and Be. D’s work was supported by Aspirant grants from the FNRS. S.R. is an FRS-FNRS research associate. This work is supported in part by the Belgian program of Interuniversity Attraction Poles initiated by the Federal Office for Scientific Technical and Cultural Affairs (PAI no. P7/44) to Ba.D. and S.R., and by the FNRS (research project T.0006.14-PDR [FRFC]) to S.R. I.M.F. was supported by the Agriculture and Food Research Initiative Competitive Grants Program (grant 2010-65110-20416 from the U.S. Department of Agriculture’s National Institute of Food and Agriculture to R. L.). Ba.D. is also supported by a BOF-basic equipment and a GOA grant from the Ghent University special research funds. The authors declare that there is no conflict of interest regarding the publication of this article.

---

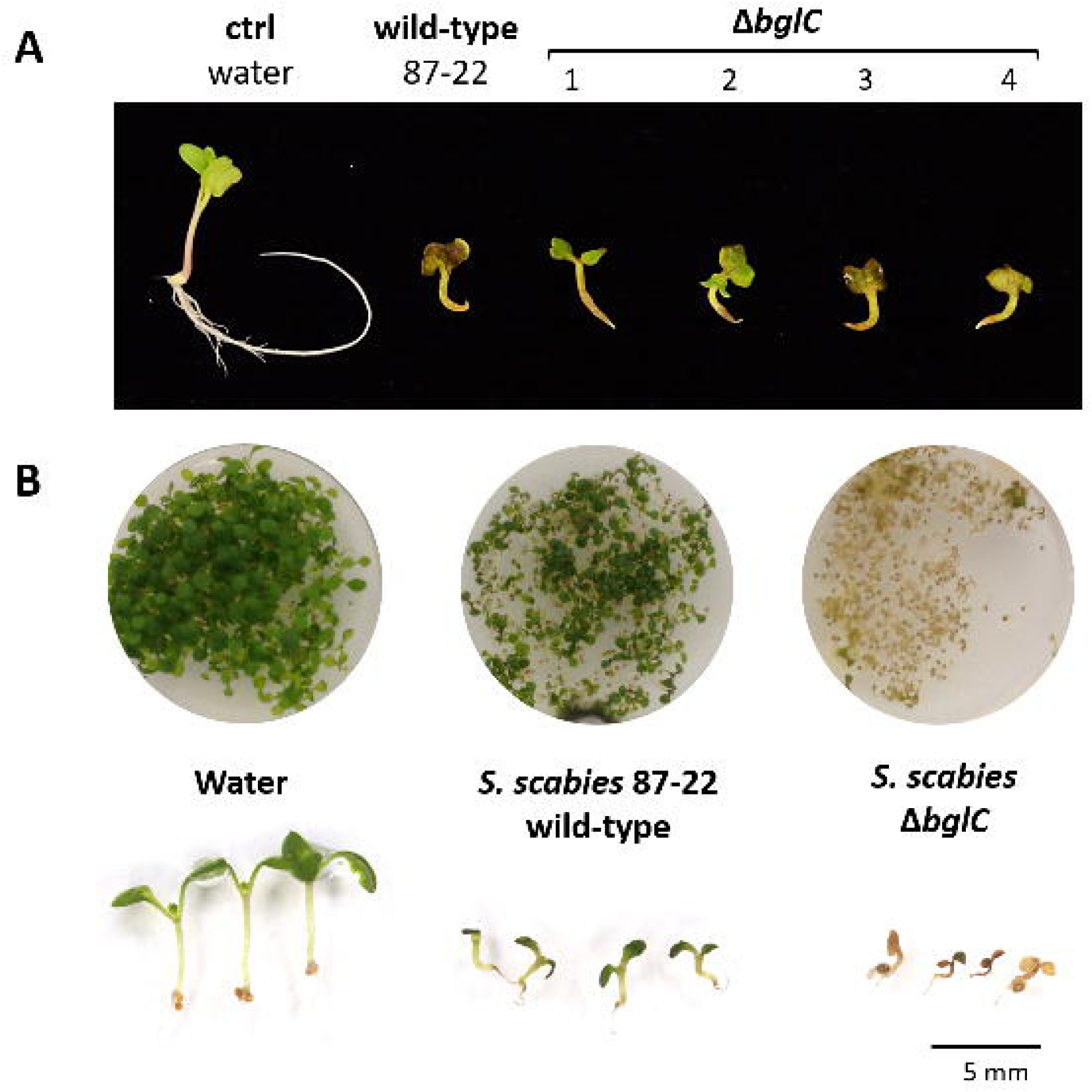

**Fig. S1.**
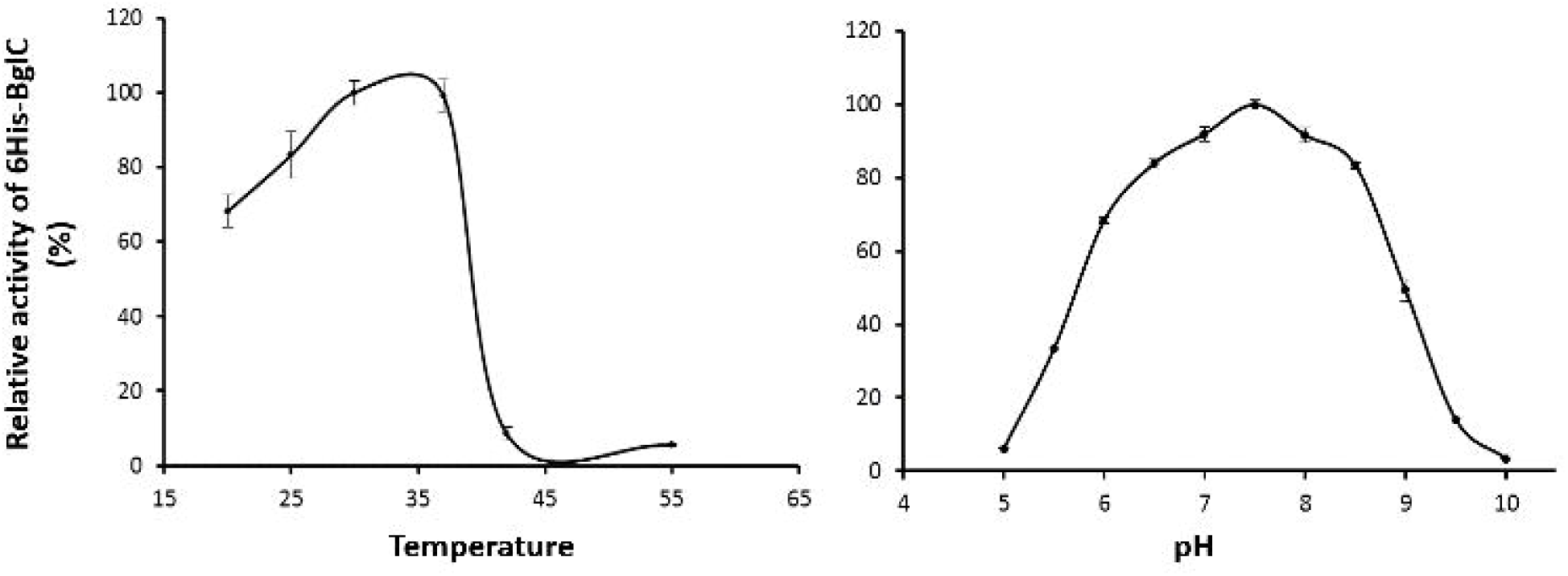
Effect of temperature and pH on BglC activity. The optimal temperature was determined by measuring the relative enzyme activity of BglC (0.2 μM) in HEPES 50 mM pH 7.5 at 20, 25, 30, 37 and 42 °C. The effect of pH on the relative activity of BglC was assessed in the range of pH 5.0 - 6.5 (50 mM MES buffer), pH 7.0-8.5 (50 mM HEPES buffer), and pH 9-10 (50 mM CHES buffer) at 25°C.

**Fig. S2.**
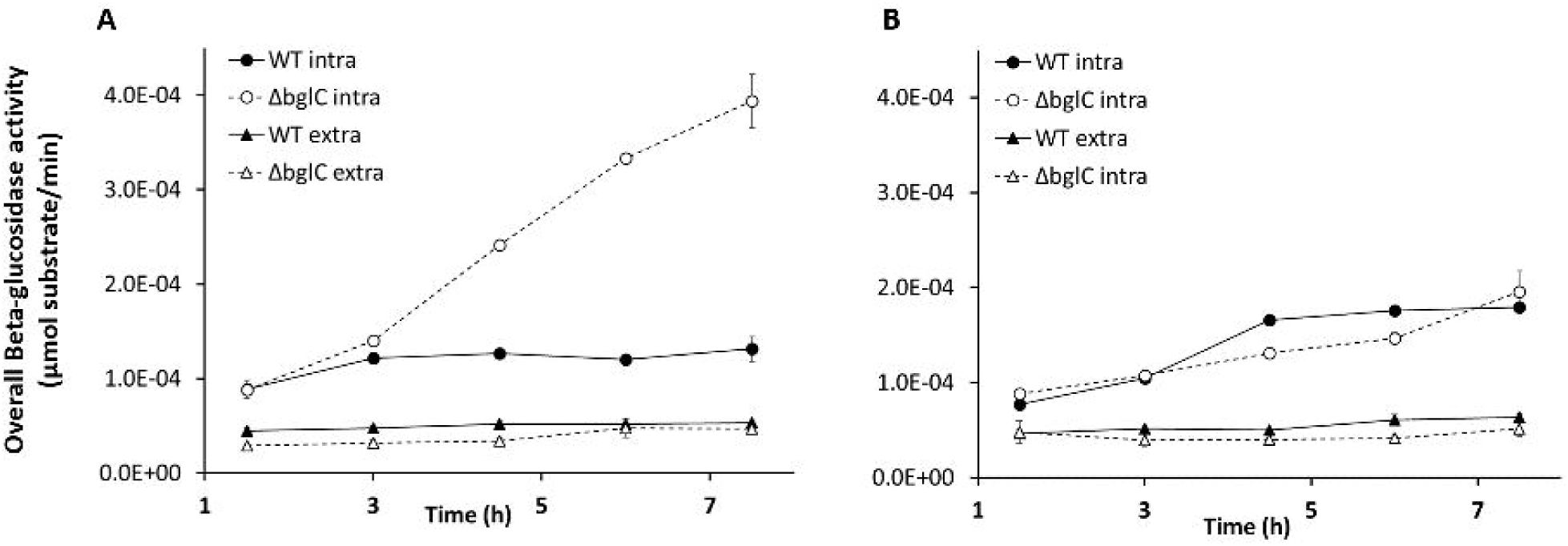
Weak overall extracellular Beta-glucosidase activity of S. scabies compared to its overall intracellular Beta-glucosidase activity. Overall intra- and extracellular β-glucosidase activity of S. scabies wild-type and its bgIC null-mutant grown in MM supplemented with cellobiose (**A**) and cellotriose (**B**).

**Fig. S3.**
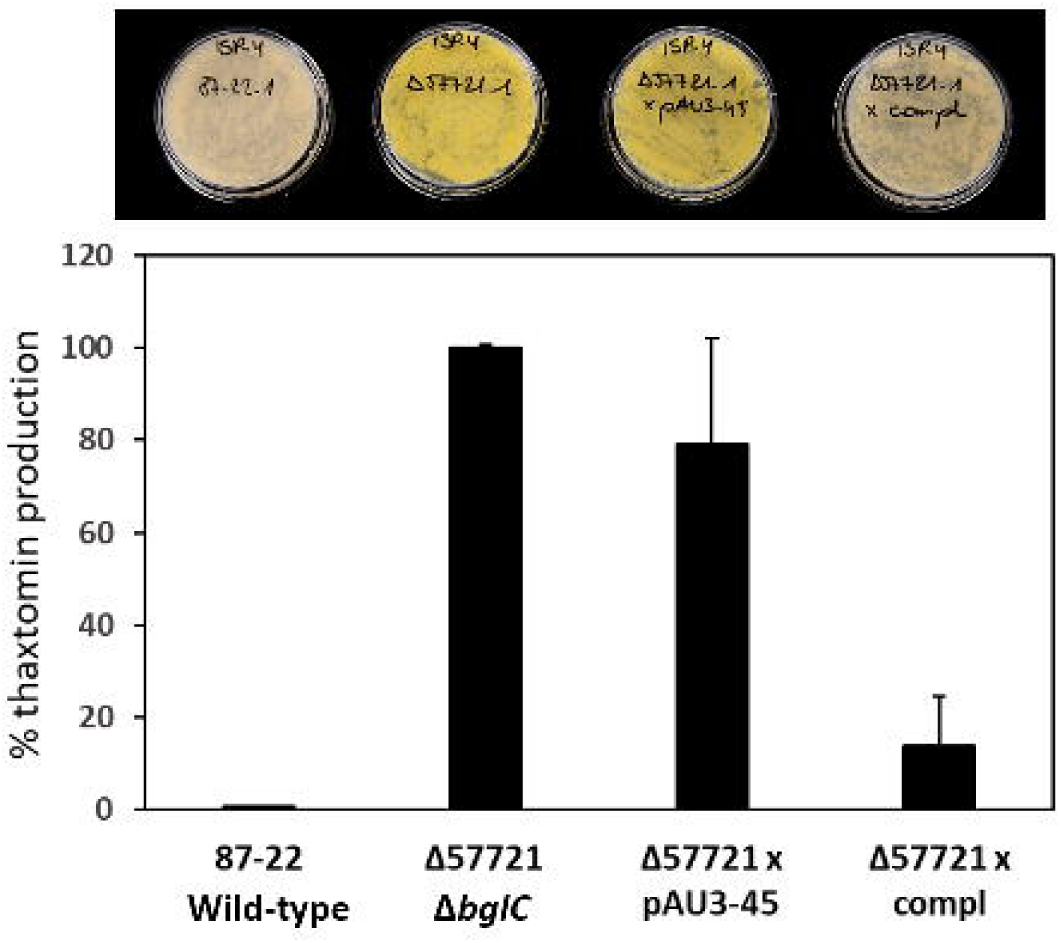
Complementation of the bglC mutant. The *bglC* mutant complemented with plasmid pIMFOOl carrying the *S. scabies bglC* gene and its upstream region restored thaxtomin production to the level produced by the wild-type, demonstrating that the phenotype of the mutant is indeed caused by the chromosomal deletion of the *S. scabies bglC* gene.

**Fig. S4.**
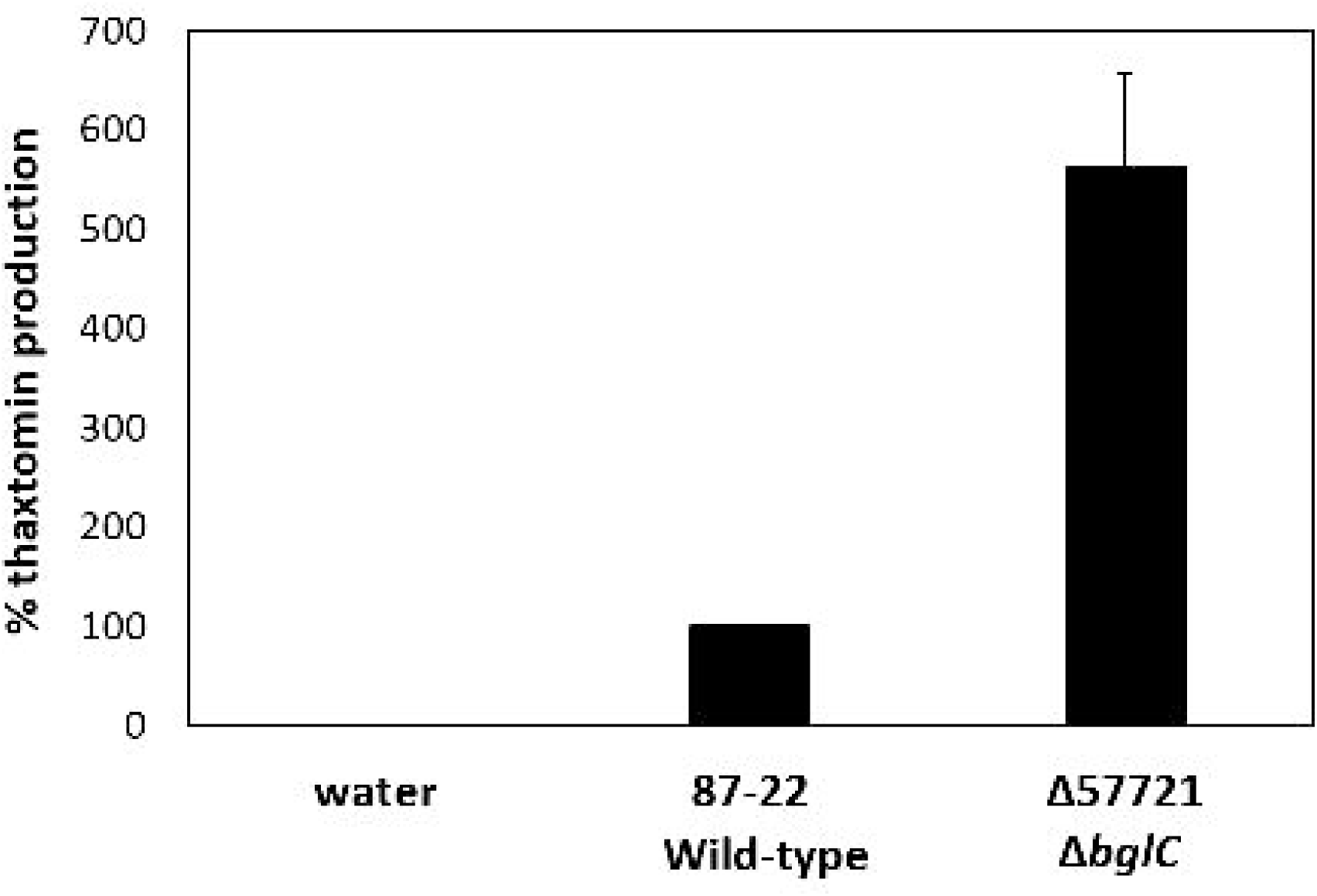
Thaxtomin A production of S. scabies wild-type the bglC mutant when inoculated on radish seedlings. HPLC analysis of thaxtomin extracted from the corresponding radish assays showing that although there is no visual difference in virulence on radish between wild-type and mutant strains (Figure 7A), the *bglC* mutant isolates produced significantly more thaxtomin than the wild-type bacteria when inoculated on plants.

